# Phosphorylation of Zearalenone retains its toxicity

**DOI:** 10.1101/2024.07.30.605906

**Authors:** Muhammad Asaduzzaman, Ivan Pavlov, Guillaume St-Jean, Yan Zhu, Mathieu Castex, Younes Chorfi, Jerome R E del Castillo, Ting Zhou, Imourana Alassane-Kpembi

## Abstract

Microbial biotransformation of Zearalenone (ZEN) is a promising deactivation approach. The residual toxicity and stability of Zearalenone-14-phosphate (ZEN-14-P) and Zearalenone-16-phosphate (ZEN-16-P), two novel microbial phosphorylation products of ZEN, remain unknown.

We investigated the cytotoxicity, oxidative stress, pro-inflammatory, and estrogenic activity of phosphorylated ZENs using porcine intestinal cells and uterine explants, and human endometrial cells, and traced their metabolic fate by LC-MS/MS analysis.

The phosphorylated ZENs significantly decreased the viability of IPEC-J2 and Ishikawa cells. Similar to ZEN, phosphorylation products induced significant oxidative stress, activated the expression of pro-inflammatory cytokines, and demonstrated estrogenic activity through upregulation of estrogen-responsive genes, activation of alkaline phosphatase and proliferation of endometrial glands. LC-MS/MS analysis pointed that although phosphorylated ZENs are partially hydrolyzed to ZEN, their respective metabolic pathways differ. We conclude that phosphorylation might not be sufficient to detoxify ZEN, leaving its cytotoxic, pro-inflammatory and estrogenic properties intact.

**Graphical abstract:** 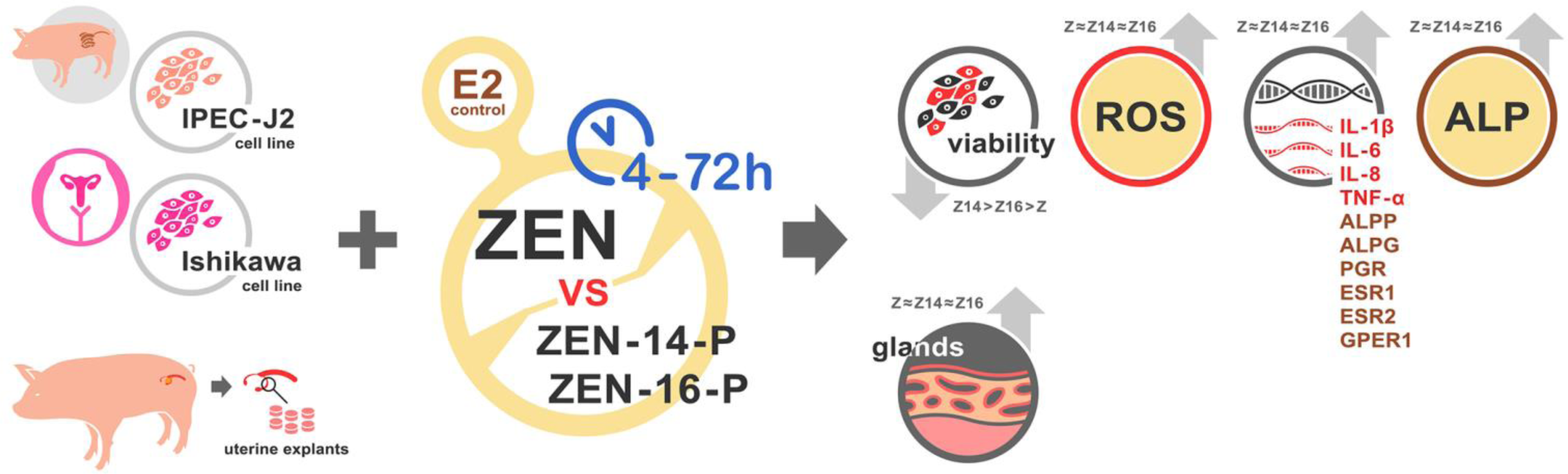

## INTRODUCTION

Fungal contamination has become a major problem around the world as it can lead to significant economic loss in global agriculture. *Fusarium* mycotoxins not only impose a serious risk to human and animal health but also lead to significant economic loss in the sector of livestock products.^1^ Zearalenone (ZEN) is one of the mycotoxins produced by *Fusarium* fungi as a secondary metabolite that plays an important role as a toxin for humans and farm animals.^2^ ZEN and its metabolites have been detected in humans and animals.^3^ Pigs, particularly prepubertal gilts, are commonly affected by ZEN.^4^ Pig feed is largely composed of grains, which represent an important source of ZEN contamination.^2^ It induces strong estrogenic effects that are attributed to the similarity between ZEN and 17-β-estradiol.^5^ ZEN is capable of binding estrogen receptors ERα and ERβ, which cause functional and morphological changes in the reproductive system.^6^ Estrogenic effects in female prepubertal gilts may exhibit symptoms such as vulva swelling and redness, ovarian cyst formation, uterine enlargement, and mammary gland enlargement. In male pigs, common manifestations include decreased sperm production and testicular atrophy.^7^ The induction of systemic inflammation by ZEN disrupts immune-related pathways that may reduce host resistance to infectious diseases.^8^ Feeding pigs ZEN-contaminated diets result in impaired immune function and decreased resistance to various infectious diseases.^9^ Moreover, ZEN induces intestinal inflammation and activates inflammatory cytokine expression.^9^ At the cellular level, ZEN induces oxidative damage of DNA owing to the accumulation of ROS.^10^

Owing to the widespread presence and adverse effects of ZEN in humans and animals, the development of effective neutralization methods is extremely important. Many decontamination strategies have been developed to reduce or eliminate ZEN such as physical, chemical, or biological methods.^11^The most common detoxification methods are physical and chemical methods. Due to the limitations of these methods associated with high costs and the loss of essential food nutrients, biological strategies are widely recognized as the most preferable methods for the removal or degradation of ZEN.^12^ A purified bacterial isolate, *Bacillus* sp. S62-W obtained from corn silage samples, exhibited the capability of transforming high concentrations of ZEN up to 200 µg/mL into phosphorylated derivatives zearalenone-14-phosphate (ZEN-14-P) and Zearalenone-16-phosphate (ZEN-16-P).^13^ This phosphorylation functionality was also detected in thirteen other *Bacillus* strains of various species, suggesting that the metabolism pathway is widely conserved in *Bacillus* spp. Although many biotransformation products of ZEN show decreased estrogenicity, some modified forms of ZEN are easily transformed into their parental form, thus contributing to ZEN exposure.^7^ As far as phosphorylated ZEN derivatives are concerned, their residual toxicity compared to ZEN and their stability are still unknown.

In this study, we investigated the hypothesis that the biotransformation of ZEN into C-14 or C-16 phosphorylation products abolishes its toxicity by inhibiting cytotoxicity, oxidative stress, pro-inflammatory activity as well as estrogenic activity, using porcine intestinal IPEC-J2 cells and uterine explants, and human endometrial cells. Since phosphorylation did not appear to decrease the toxicity of ZEN, we traced the metabolic fate of the phosphorylated ZEN derivatives by LC-MS/MS analysis, to clarify their intrinsic toxicity.

## MATERIALS AND METHODS

### Chemicals and Reagents

ZEN-14-P and ZEN-16-P used in this study were obtained as gifts from Prof. Dr. Ting Zhou (Agriculture and Agri-Food Canada). ZEN was purchased from Fermentek (Fermetek Ltd, Jerusalem, Israel). ZEN-14-P, ZEN-16-P, and ZEN dissolved in dimethylsulfoxide (DMSO) at 25 mM were stored at −20°C before dilution in a culture medium. The DMSO used in experiments was purchased from Sigma-Aldrich (Oakville, ON, Canada). Cell culture medium, PBS solution (PBS), antibiotics (penicillin and streptomycin), fetal bovine serum (FBS), trypsin, insulin-transferrin-selenium (ITS), epidermal growth factor (EGF) were purchased from Wisent (Wisent Bioproducts, St-Bruno, QC, Canada). Thiazolyl blue tetrazolium bromide (MTT) and stop/solubilization solution from Promega (Promega, Madison, WI) were used in the MTT test. 2′,7′-dichlorodihydrofluorescein diacetate (DCF-DA) from Abcam (Abcam, Waltham, MA, USA) was used in oxidative stress determination. 17-β-estradiol, *P*-nitrophenylphosphate, MgCl_2_, diethanolamine, and estrogen receptor antagonist (ICI 182 780) used in alkaline phosphatase activity tests were purchased from Sigma-Aldrich (Sigma-Aldrich, Oakville, ON, Canada). PureZOL^TM^ RNA purification reagent from BIO-RAD (BIO-RAD, California, USA), Advanced cDNA synthesis kit and qPCR Master mix from Wisent (Wisent Bioproducts, St-Bruno, QC, Canada) were applied in real-time PCR analysis. Formaldehyde: Tissuefix (Chaptec, Montreal, QC, Canada), Hematoxylin from Fisher Scientific (Fisher Scientific, Hampton, New Hampshire, USA), and Saffron from Chaptec (Chaptec, Montreal, QC, Canada) were used to assess the proliferation of porcine endometrial glands.

### Cell culture

Ishikawa human adenocarcinoma cell line from ATCC and intestinal porcine jejunal epithelial cell line (IPEC-J2) from DSMZ were maintained in DMEM-F12 medium supplemented with insulin-transferrin-selenium (ITS), fetal bovine serum (FBS), epidermal growth factor (EGF) and antibiotics (penicillin and streptomycin). The cells were maintained at 37°C in a humidified atmosphere containing 5% CO_2_ in a cell incubator.

### Assessment of Cell Viability by MTT Assay

IPEC-J2 and Ishikawa cells were seeded in 96-well tissue culture plates (10^4^ cells /well in 100 µL medium) (Ultident, St-Laurent, QC, Canada). The cells were cultured in an incubator at 37°C and 5% CO_2_ for 24 h, after that the cell culture medium was replaced with FBS-free medium containing graded concentrations of mycotoxins ZEN-14-P, ZEN-16-P, and ZEN. The tested concentrations of compounds ranged from 0.78 μM to 100 μM. After 48 h or 72 h of mycotoxin exposure, 15 μl MTT solution was added in each well and the cells were cultured for another 4 h at 37°C. Finally, 100 μl stop/solubilization solution was added in each well in this experiment, and after an additional hour room temperature incubation, absorbance at 570 nm was recorded using a SpectraMax i3 microplate reader (Molecular Devices, San Jose, CA). Calculation of percentage of cell viability was achieved according to the equation:

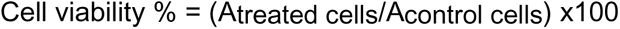

Where, Atreated cells is the optical density in the treated wells, and Acontrol cells is the optical density of the control wells.

For each toxin, the individual cell viability vs. concentration data sets were subjected simultaneously to nonlinear mixed effect modeling in ADAPT, version 5.^14^ The parameters I_max_, IC_50_ and H of the inhibitory sigmoid E_max_ model

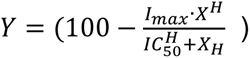

were estimated using MLEM methods, under the assumption that they are uncorrelated and log-normally distributed. These parameter estimates, which respectively describe the efficacy, potency and steepness of the relationship between the percent viability (Y) and toxin concentration (X), were retained for statistical analysis.

### Assessment of Oxidative Stress

Intracellular ROS levels were detected using a specific fluorescent probe dye DCF-DA according to the manufacturer’s instructions. Briefly, 2×10^5^ IPEC-J2 cells/well in 1 mL medium were seeded in 24-well tissue culture plates (Ultident, St-Laurent, QC, Canada) and allowed to adhere for 24 h. The cell culture medium was replaced with an FBS-free medium containing 10 μM test compounds for 4 h. The cells were then stained with DCF-DA for 45 mins at 37°C in the dark, and washed with buffer to remove any unincorporated dye. The green fluorescence intensity was measured using a fluorescence microscope (Axio Imager M1, Zeiss, Germany). The data were analyzed using the ImageJ software (NIH, Bethesda, MD, USA).

### Assessment of Alkaline Phosphatase Activity

Ishikawa cells were plated at a density of 10^4^ cells/well in 100 µL phenol-red free medium supplemented with charcoal-stripped FBS and seeded in 96-well flat-bottom tissue culture plates (Ultident, St-Laurent, QC, Canada) for 24 h. The medium was replaced by 100 µL FBS-free medium and allowed to settle for the next 24 h, after which the cell medium was replaced with FBS-free medium containing different concentrations of tested compounds (ZEN-14-P, ZEN-16-P, ZEN and 17-β-estradiol) for the next 48 h. After dilution, the concentrations of compounds ranged from 10^−6^ nM to 10^−2^ nM. An estrogen receptor antagonist (ICI 182 780) was used to verify whether the detected effects were estrogen dependent. Each compound’s concentration was treated with 1 µM estrogen receptor antagonist (ICI 182 780). After 48 h, the medium was removed and the plates were rinsed twice with 200 μl PBS. Afterwards, the plates were placed at −80°C for 30 min followed by 5 min thawing at room temperature. The plates were then placed on ice and 50 μl ice-cold solution (pH 9.8) containing 5 mM *p-*nitrophenyl phosphate, 0.24 mM MgCl_2,_ and 1M diethanolamine was added. Plates were then read periodically at 405 nm light wavelength in SpectraMax i3 plate reader (Molecular Devices, San Jose, CA) for 1 h at 3 min intervals. The ALP activity was calculated as the slope (unit/min) of the curve.

Due to the presence of outlier values, time vs. optical density (OD) relationships were subjected to robust linear regression in SAS, using the MM estimation method.^15^ After visual verification of the goodness of fit of the resulting predictions, the estimated robust slopes and the raw OD values measured at 60 minutes (final reading time) were retained for statistical analysis.

### Total RNA Extraction and Real-time PCR

Expression of the estrogen-responsive genes *ALPG*, *ALPP*, *PGR*, *ESR1*, *ESR2* and *GPER1* in Ishikawa cells and pro-inflammatory cytokine genes *IL-1β*, *IL-6*, *IL-8* and *TNF-α* IPEC-J2 cells following exposure to mycotoxins was analysed using specific primers (Table 1). Ishikawa cells were seeded at 5×10^5^ cells per well in 3 mL phenol-red free medium supplemented with charcoal-stripped FBS in six-well plates (Ultident, St-Laurent, QC, Canada) for 48 h. After 24 h exposure to 10 μM ZEN-14-P, or ZEN-16-P, or ZEN, the medium was removed and total RNA was extracted using PureZOL^TM^ RNA purification reagent according to the manufacturer’s instructions. To analyze pro-inflammatory cytokine gene expression, IPEC-J2 cells were seeded in 6-well tissue culture plates (5×10^5^ cells/well in 3 mL of medium) and allowed to adhere for 24 h. Cells were re-fed with media three times weekly for 14 days and used after reaching confluency. Cells were then rinsed with PBS and treated for 48 h with FBS-free media containing 5 or 25 μM of each tested compound. After 48 h of mycotoxin exposure, total RNA was extracted using PureZOL^TM^ RNA purification reagent according to the manufacturer’s instructions. After checking the purity and concentration using a Nanodrop spectrophotometer, reverse transcription was performed using an advanced cDNA synthesis kit according to the manufacturer’s protocol. Real-time PCR was performed on a CFX96 Touch Real-Time PCR detection system (Bio-Rad) using qPCR Mastermix following the manufacturer’s protocol. All primers were tested to achieve amplification efficiencies between 90 % and 110 %. Samples were run in duplicates, and data were expressed relative to *GAPDH* and *RPL-32* used as housekeeping genes to analyze the expression of estrogen-responsive and pro-inflammatory cytokine genes, respectively and normalized to calibrator sample using the Pfaffl method ^16^ with correction for amplification efficiency.

**Table 1.**
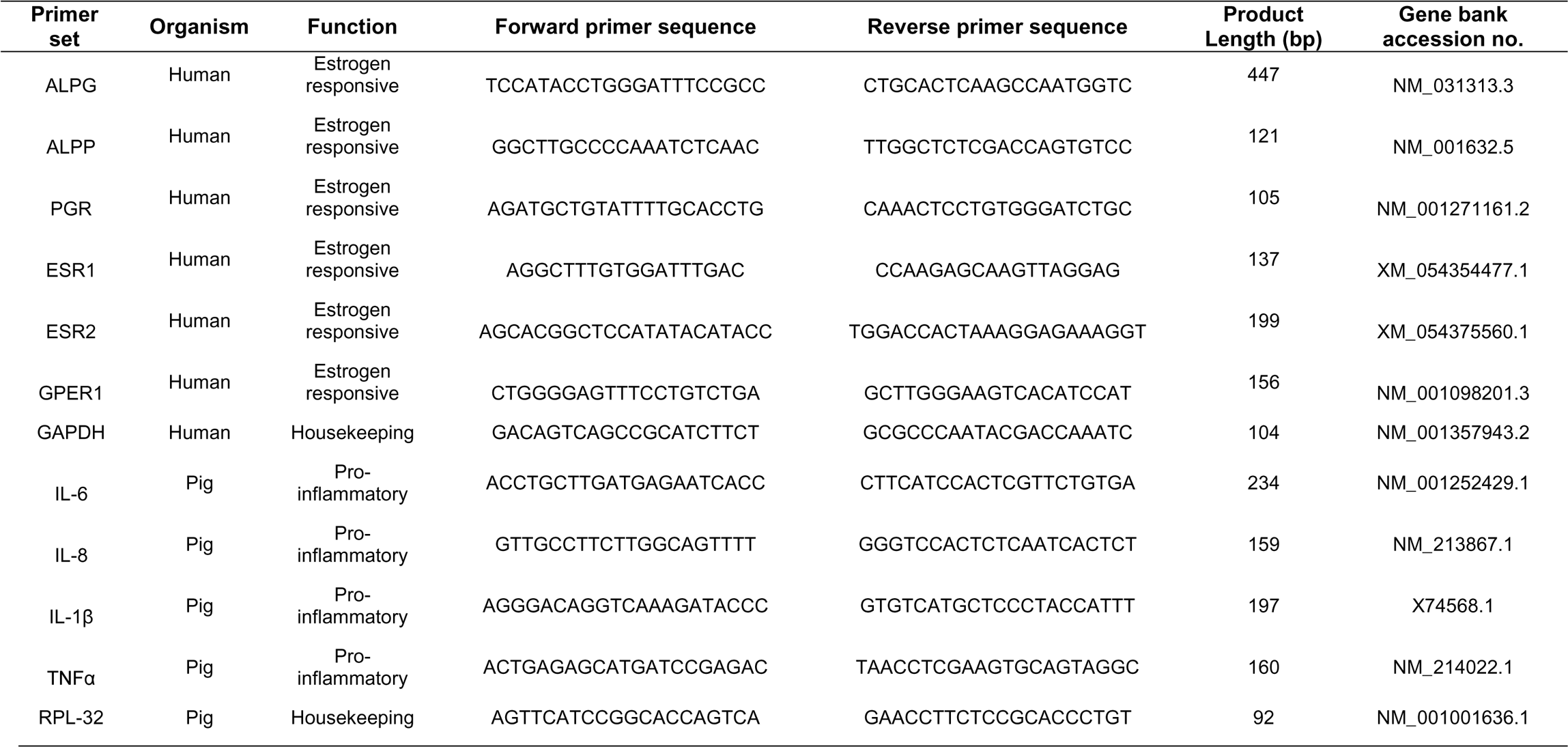
Primer sequences for the estrogen-responsive and pro-inflammatory cytokine genes used in this study.

### Assessment of the Proliferation of Porcine Endometrial Glands

The experiment was conducted according to the Guide to the Care and Use of Experimental Animals published by the Canadian Council on Animal Care. The Ethics committee of the University of Montreal “Comité d’Éthique de l’Utilisation des Animaux de l’Université de Montréal” approved all animal experimentation procedures (#Rech-2071).

Uteri were collected from pre-pubertal gilts (n=6), <100 days old. To prepare for endometrial explants, the uterine horns were cut into shorter segments. Endometrial explants were punched out from the segments using a 6mm biopsy punch tool (Integra Bioscences, Hudson, NH, USA) and pre-incubated in a petri dish in a warm (37° C) culture medium. Explants were exposed to 30 μM tested compounds in 6-well flat-bottom tissue culture plates (Ultident, St-Laurent, QC, Canada) for 24 h. After those explants were fixed in 10% formalin and stained with Hematoxylin phloxine saffron (HPS), slides were scanned with Aperio AT2 scanner, and evaluated with Aperio ImageScope V12.4.3 (Leica Biosystems, Wetzlar, Germany).

### Investigation of Metabolic fate of ZEN and Phosphorylated ZENs

IPEC-J2 cells at a density of 5×10^5^ cells/well in 3 mL were seeded in six-well tissue culture plates (Ultident, St-Laurent, QC, Canada) and allowed to adhere for 24 h. Cells were re-fed with media three times weekly after reaching confluency and used 14 days after seeding. Ishikawa cells were seeded at 5×10^5^ cells/well in 3 mL of phenol-red free medium supplemented with charcoal-stripped FBS in six-well plates (Ultident, St-Laurent, QC, Canada) for 48 h. Both cell types were then rinsed with PBS and treated for 24 h with FBS-free media containing 25 μM of each of the tested compounds. Cell culture supernatant samples were purified using Phenomenex Strata X C18 solid phase extraction cartridges (30 mg) and eluted with 1 % (v/v) formic acid in methanol. LC-MS/MS analysis was performed using a Thermo® Scientific Q-Exactive™ Orbitrap mass spectrometer equipped with a Vanquish™ Flex Binary UPLC System (Waltham, MA, USA). A Kinetex XB-C18 100A column (100 x 4.6 mm, 2.6 µm, Phenomenex Inc., Torrance, CA, USA) was used. The binary mobile phase consisted of solvent A (99.9% H_2_O / 0.1% formic acid) and solvent B (99.9% AcN / 0.1% formic acid). The solvent gradient was 0-20 min, 20% to 80% B; 20-21 min, 80% to 100% B; 21-24 min, 100% B; 24-25 min, 100% to 20% B; 25-31 min, 20% B. The column temperature was set to 23°C, the flow rate was set to 0.700 mL/min, and the injection volumes were 2 uL, 4 uL and 10 µL. UV peaks were monitored at 275 nm. Both positive and negative ionization modes were used for ion detection, with the spray voltage set to 4.75 kV. Mass spectrometry data were collected using the Full-MS/DDMS2 (TopN = 10) method, with NCE set at 30 and intensity threshold set at 1.0e5 counts. Data were visualized and analyzed using Thermo FreeStyle™ 1.7PS2 software (Thermo Fisher Scientific, Mississauga, ON, Canada).

### Statistical analysis

All statistical analyses were performed using generalized linear mixed models,^17^ in SAS, 9.4 version (SAS Institute, Cary NC, U.S.A.). Except for endometrial proliferation, which is a discrete count variable, all other variables were continuously distributed. The most suitable statistical distribution was selected for each response variable based on the Akaike information criterion of goodness of fit.^18^ Exploratory data analyses were performed to identify the potentially relevant fixed-effect factors (e.g., toxins, concentrations, exposure times, cell lines, etc.) and interactions to include in each model, and the study replicate was used as a random effect in most models.

Two families of *a priori* simultaneous unilateral tests were performed using the estimated least-square means of the models. The first, upper-tailed test, assessed the toxicities of the tested substances as compared to the negative control. The second, lower-tailed test, assessed the detoxication efficiency of ZEN-14-P and ZEN-16-P relative to ZEN. Because the two families of tests had opposite directions (lower vs. upper tail), the overall 0.05 alpha level was split proportionately to the number of pairwise comparisons of reference vs. test level to perform in each family. To control the probability of erroneously rejecting one or more hypotheses in the resulting series of statistical tests while preserving its statistical power, the stepdown hybrid Monte-Carlo method of Westfall (1997) ^19^ was used.

## RESULTS

### C-14 or C-16 phosphorylation increases the cytotoxicity of ZEN

To know if the biotransformation of ZEN by *Bacillus sp* S62-W affects it cytotoxic effect, we compared the effects of ZEN, ZEN-14-P and ZEN-16-P on the viability of the porcine intestinal IPEC-J2 cells and the human endometrial Ishikawa cells, after 48 h and 72 h of exposure. Figure 1 and Table 2 present the parameters for the dose-response relationship of the cytotoxic effects observed for the different compounds. In IPEC-J2 cells, both phosphorylated derivatives were more potent cell death inducers. On the other hand, both the I_max_ and IC_50_ parameters in Ishikawa cells consistently indicated that C14 or C16 phosphorylation does not decrease the cytotoxicity of ZEN, but rather increase it.

**Figure 1.**
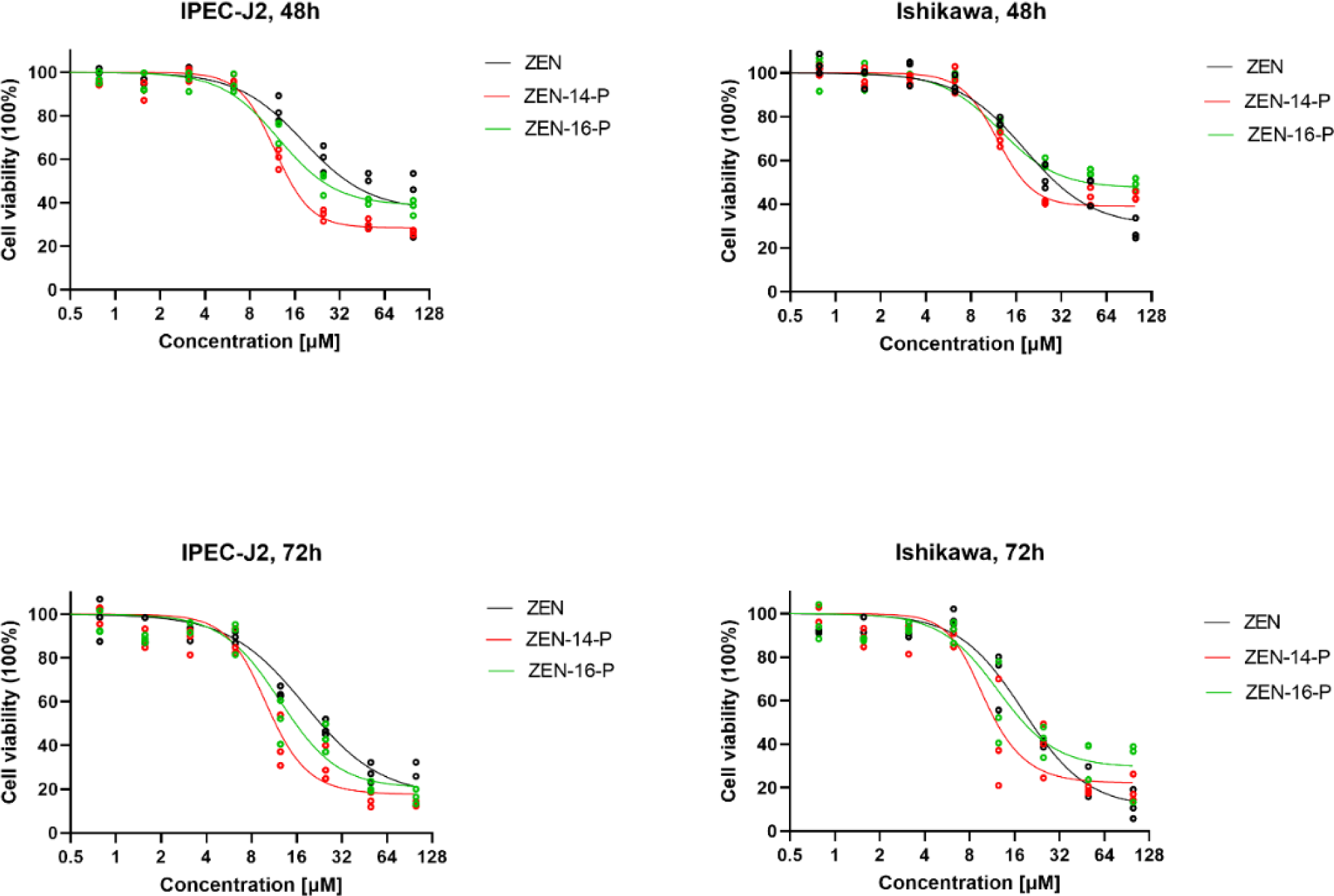
Comparative cytotoxic effect of ZEN and C-14 and C-16 phosphorylation of ZENs. Ishikawa and IPEC-J2 cells were exposed for 48 h or 72 h to ZEN, ZEN-14-P or ZEN-16-P. Cytotoxicity was assessed by the MTT (Dots: % viability values of three independent replicates; lines: mean of individual conditional values predicted by the sigmoid inhibitory Emax model)

**Table 2.**
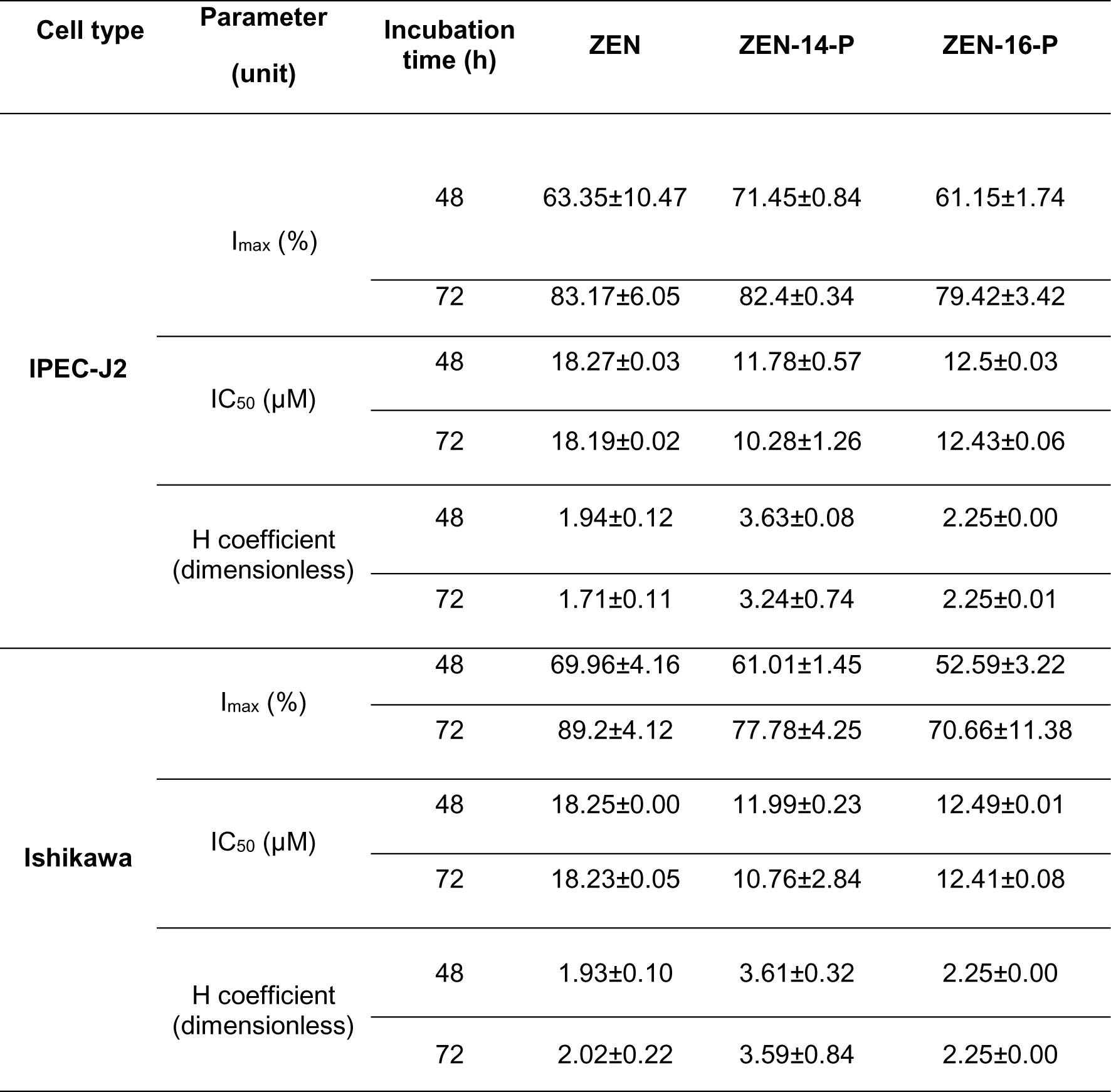
Efficacy (Imax), Potency (IC_50_) and Steepness (H) parameters of the cytotoxicity assay, as estimated with the nonlinear mixed-effect model using the MLEM estimation methods.

### C-14 or C-16 phosphorylation does not prevent oxidative stress induction by ZEN

Excessive ROS can cause cellular oxidative stress. To further explore whether phosphorylation affects ROS induction by ZEN, we analyzed ROS levels after treatment of IPEC-J2 cells with 10 μM of either ZEN, or ZEN-14-P, or ZEN-16-P for 4 h. Cell fluorescence indicating the presence of ROS significantly differed across incubation conditions (p < 0.0001) (Figure 2A and 2B). Compared with the negative control, exposure to 10 μM ZEN-14-P, ZEN-16-P and ZEN significantly increased cell fluorescence. Yet, both ZEN-14-P and ZEN-16-P induced as much ROS as ZEN, and hence the unilateral tests targeting a detoxifying effect were not significant (p = 1.0).

**Figure 2.**
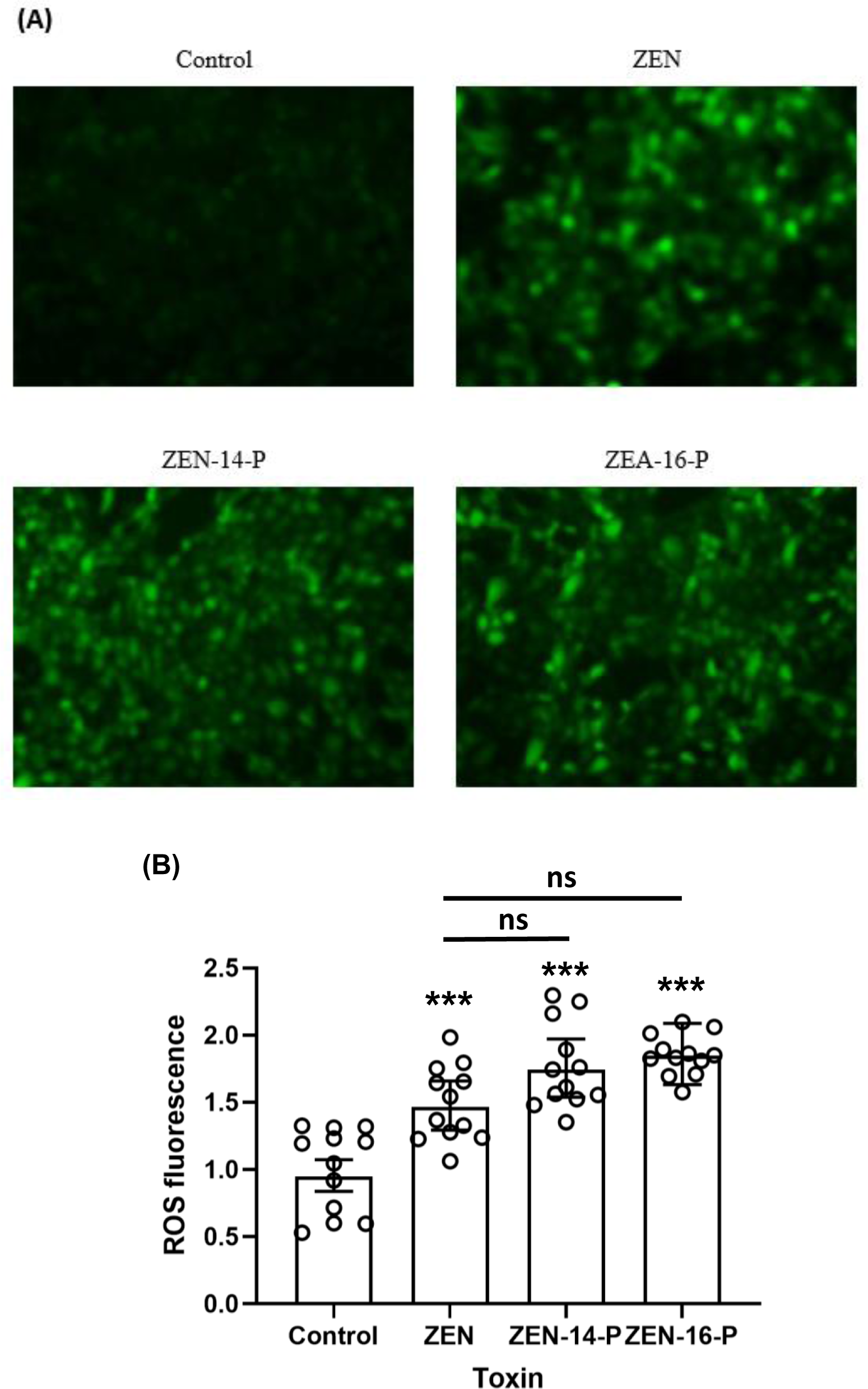
Comparative oxidative stress induction of ZEN and of C-14 and C-16 phosphorylated ZENs. IPEC-J2 cells were exposed to 10 µM ZEN, ZEN-14-P or ZEN-16-P for 4 h. (A) ROS-induced fluorescence of cells exposed to the tested compounds under the fluorescence microscope (B) Green fluorescence intensity values associated with the ROS production of IPEC-J2 cells exposed to ZEN and C-14 and C-16 phosphorylation of ZENs. Dots: measured fluorescence in four separated fields of view in three independent replicates; Bars and whiskers: least-square means (LSM) and 95% confidence interval (CI). Compared to the negative control, ***p ≤ 0.001. Compared with ZEN, ns, not significant.

### C-14 or C-16 phosphorylation does not alter the pro-inflammatory activity of ZEN

We compared *in vitro* the effects of ZEN, ZEN-14-P and ZEN-16-P on the expression of pro-inflammatory cytokines. The expression of selected cytokine-encoding genes *IL-1β, IL-6*, *IL-8*, and *TNF-α* was analyzed in IPEC-J2 cells following exposure to 5 μM or 25 μM of ZEN or ZEN-14-P, or ZEN-16-P for 48 h (Figure 3).

**Figure 3.**
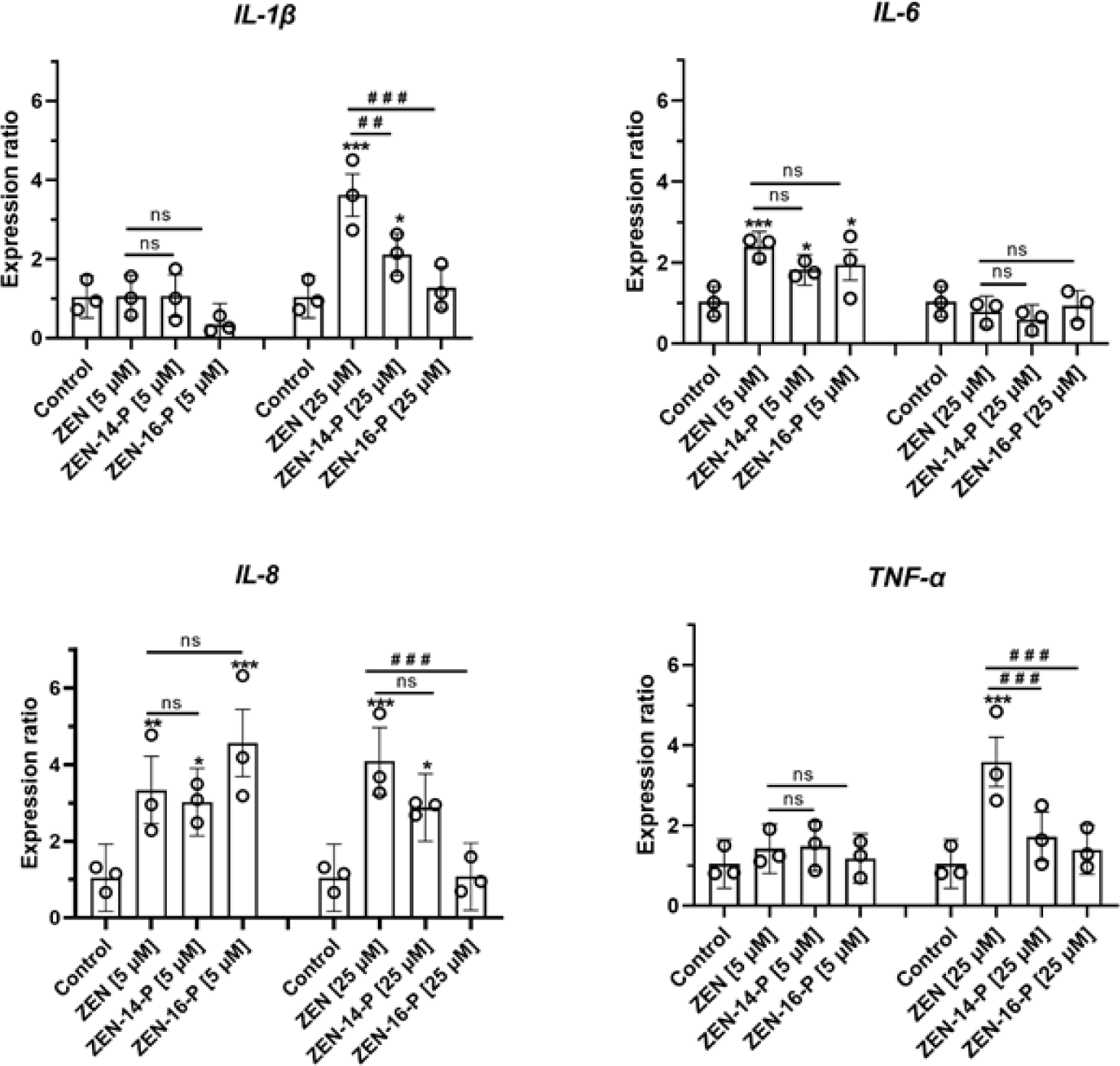
Relative expression of proinflammatory cytokine genes (*IL-6*, *IL-8*, *TNF-α*, *IL-1β*) in IPEC-J2 cells treated with 5 µM or 25 µM ZEN, ZEN-14-P, ZEN-16-P for 48 h. (Circles: measured values of three independent replicates; bars and whiskers: least-square means (LSM) and 95% confidence interval (CI)). Compared to the negative control, *p ≤ 0.05; **p ≤ 0.01; ***p ≤ 0.001; ns, not significant. Compared with ZEN, ##p ≤ 0.01, ###p ≤ 0.001; ns, not significant.

Compared to the negative control, ZEN, ZEN-14-P and ZEN-16-P showed no effect on IL-1β at 5 μM, but activated the expression at 25 μM. On the other hand, exposure to 5 μM ZEN, ZEN-14-P, and ZEN-16-P significantly upregulated the expression of IL-6, while no pro-inflammatory effect appeared at 25 μM. ZEN, and phosphorylated ZENs as well, significantly upregulated the expression of IL-8 at 5 μM. Only the exposure to 25 μM ZEN and ZEN-14-P significantly upregulated IL-8. Finally, ZEN and phosphorylated ZEN activated the expression of TNF-α at both 5 and 25 μM.

Overall, the unilateral lower-tailed tests indicate that the pro-inflammatory activity of ZEN at 5µM was not impeded by C-14 or C-16 phosphorylation. The detoxifying effects of ZEN phosphorylation were significant only at exposures of 25 µM: both ZEN-14-P and ZEN-16-P decreased the expression ratios of IL-1β (p ≤ 0.001 for both toxins) and TNF-α (p ≤ 0.0004 for both toxins). In addition, only C-16 phosphorylation of ZEN decreased the expression ratio of IL-8 (p < 0.0001).

### C-14 or C-16 phosphorylation does not decrease the estrogenicity of ZEN

The effects of C-14 or C-16 phosphorylation on the estrogenic activity of ZEN were assessed *in vitro,* using the human endometrial Ishikawa cells, and *ex-vivo,* using endometrial explants of prepubertal gilts.

First, the estrogen responsive alkaline phosphatase (ALP) activity on one hand, and the expression of estrogen responsive genes, on the other hand, were explored in endometrial Ishikawa cells exposed to either ZEN or the phosphorylated ZENs (Figure 4). Figure 4A shows the least-square mean values of the slopes of the measurements over 1 h of the optical densities of cell lysates incubated with alkaline phosphatase buffer. The activity of ALP in all the mycotoxin conditions was determined in comparison with 17-β-estradiol, after 48 h exposure of the endometrial cells with concentrations ranging from 10^−6^ nM to 10^−2^ nM. All tested substances demonstrated a dose-dependent induction of ALP activity that was suppressed in the presence of the ER high-affinity antagonist ICI 182 780, thus underlining the ER dependency of the observed ALP activity induction.

**Figure 4.**
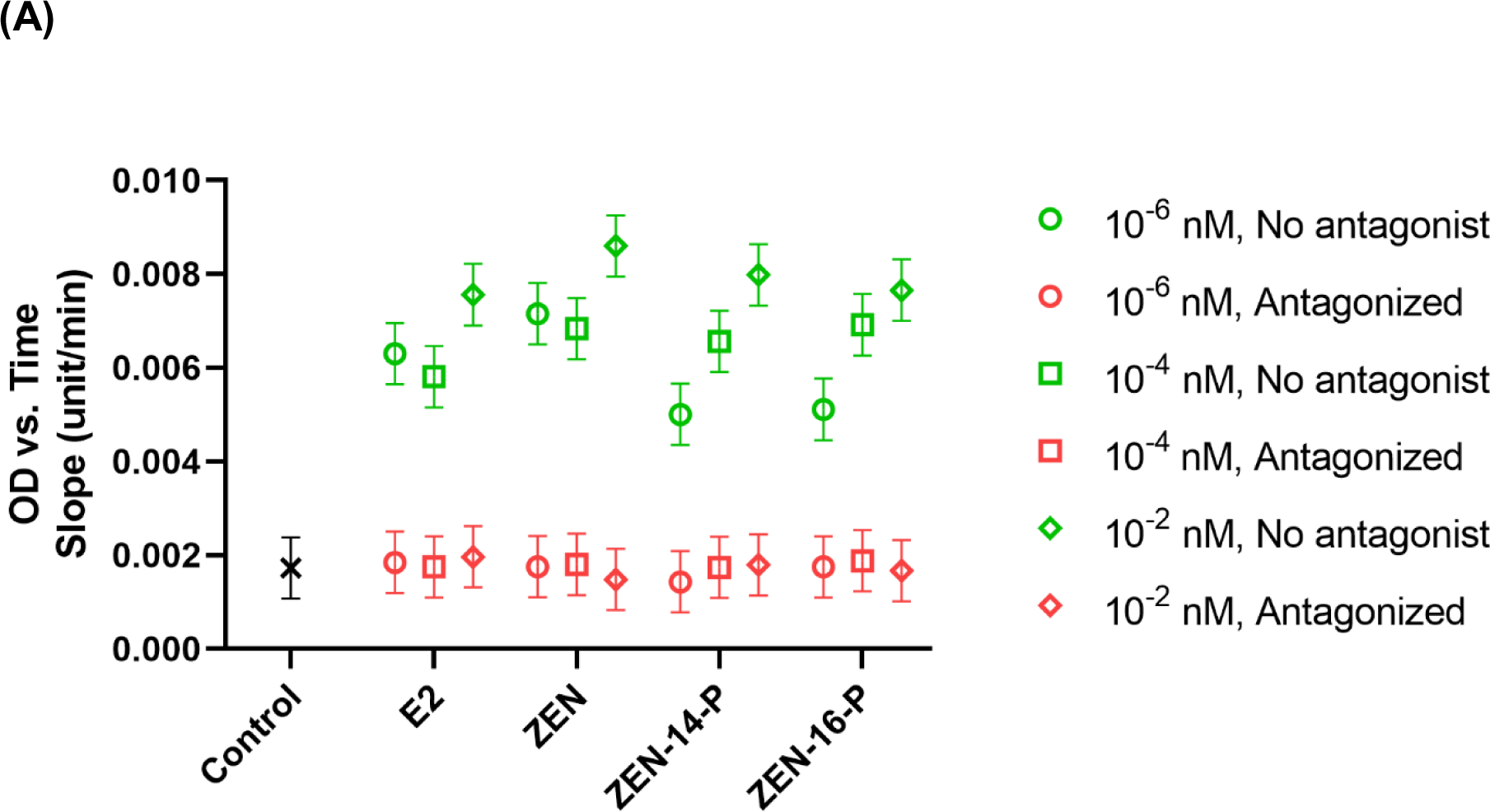
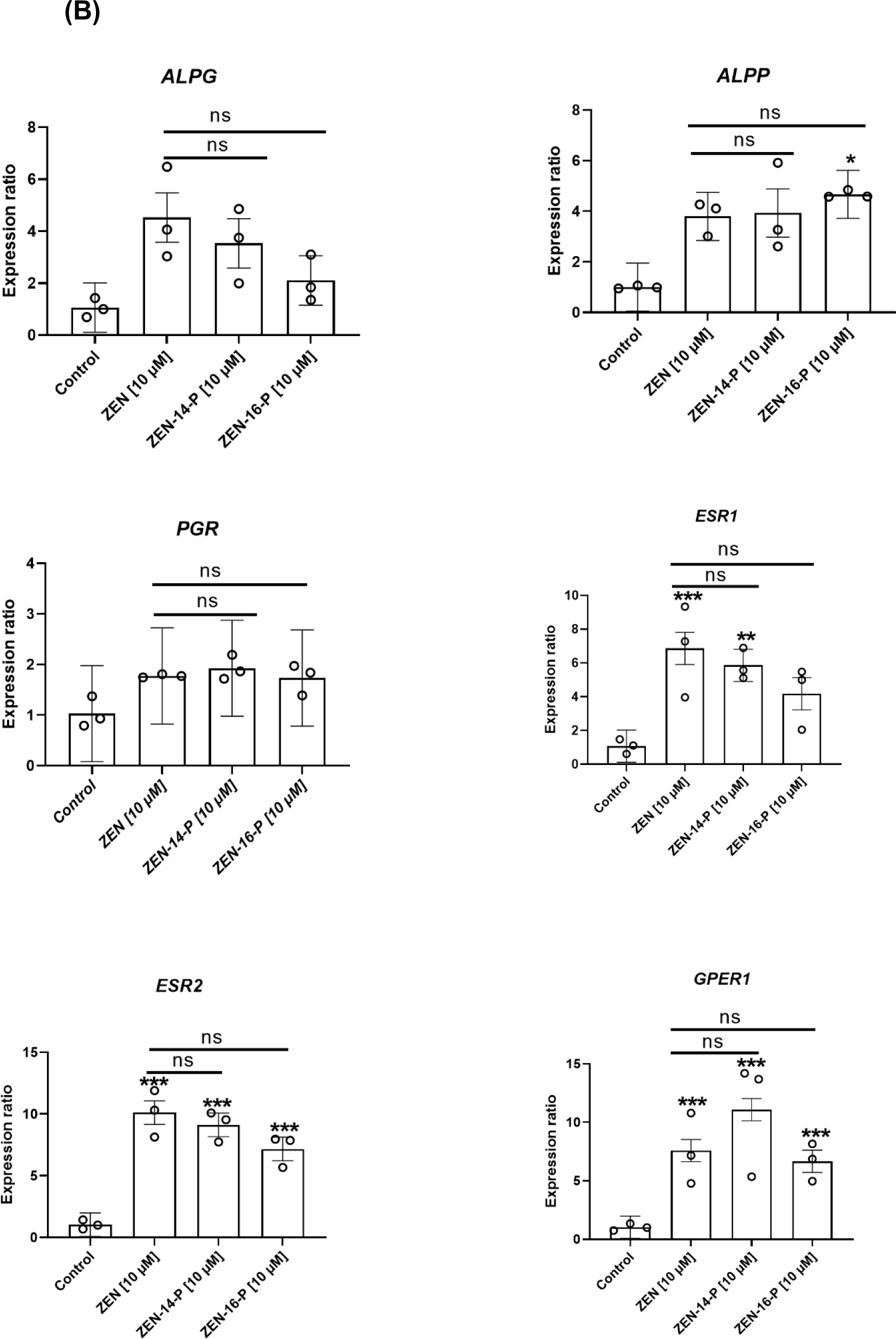
Comparative alkaline phosphatase activity and estrogen-responsive gene activation by ZEN and phosphorylated ZENs. (A) Comparative alkaline phosphatase (ALP) induction of ZEN and phosphorylated ZENs using Ishikawa cells. Ishikawa cells were exposed for 48 h to 17-β-estradiol. ALP activity was calculated as the slope of the curves, obtained by the measurements over 1 h of the optical densities of cell lysates incubated with alkaline phosphatase buffer. The concentrations of compounds ranged from 10^−6^ nM to 10^−2^ nM. The ALP activity of all tested substances was suppressed in the presence of the ER high-affinity antagonist ICI 182 780. The reported values are the least square of means (LSM) ± standard error of the mean (SEM) of at least three independent replicates. (B) Relative expression of the estrogen responsive genes *ALPG*, *ALPP*, *PGR*, *ESR1*, *ESR2*, *GPER1* in Ishikawa cells treated with 10 µM ZEN, ZEN-14-P, ZEN-16-P for 48 h. The reported values are the least-square means (LSM) ± standard error of the mean (SEM) of at least three independent replicates. Compared with the negative control, *p ≤ 0.05; **p ≤ 0.01; ***p ≤ 0.001; ns, not significant. Compared with ZEN, ns, not significant.

The series of unilateral tests of the hypothesized detoxifying effect of ZEN phosphorylation yielded non-significant reductions in slope for both ZEN-14-P and ZEN-16-P, irrespective of concentration (p < 0.10).

Figure 4B compares the expression levels of two alkaline phosphatase isoenzyme genes and selected upstream estrogen-responsive genes in Ishikawa cells exposed to either ZEN, or ZEN-14-P or ZEN-16-P on estrogen-responsive genes. The expression levels of the placental alkaline phosphatase gene *ALPP*, the germ cell alkaline phosphatase gene *ALPG*, the progesterone receptor gene *PGR*, the estrogen receptor genes *ESR1* and *ESR2*, and the G protein-coupled estrogen receptor gene *GPER1* increased dramatically upon 24 hour-exposure of Ishikawa cells to 10 µM of ZEN, or ZEN-14-P, or ZEN-16-P.

Noteworthy, the activation ratios of *ALPG*, *ESR1* and *ESR2* were lower with ZEN-16-P than ZEN, but none of these reductions were large enough to be significant in the unilateral statistical testing (p > 0.16). Overall, there was no difference in the ALP isoenzyme gene activation by ZEN or phosphorylated compounds, nor in their upstream gene regulation.

Figure 5 presents the comparative estrogenicity of ZEN and phosphorylated ZENs and the positive control 17-β estradiol (E2), as assessed *ex-vivo* in gilts uterine explants. Compared to the negative control, exposure to 30 μM E2, or 30 μM of ZEN or ZEN-14-P, or ZEN-16-P induced significant proliferation of endometrial glands in the uterine explants (p < 0.0027 for E2; p < 0.0016 for ZEN and ZEN-16-P; p < 0.021 for ZEN-14-P). Only ZEN-14-P induced less proliferation of endometrial glands than ZEN, but the reduction was not large enough to reject the null hypothesis of no detoxifying effect.

**Figure 5.**
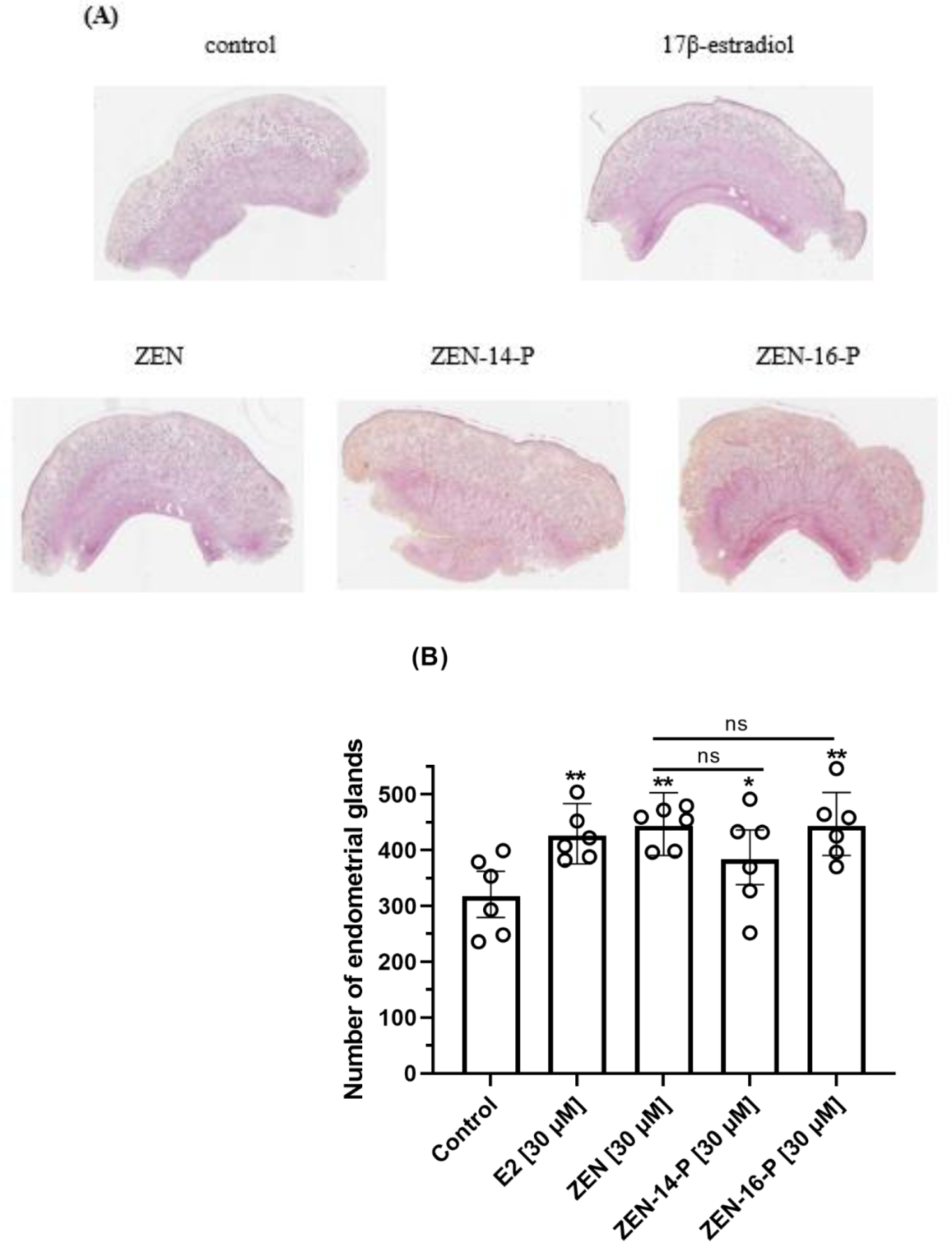
Comparative estrogenic effect of ZEN and C-14 and C-16 phosphorylation of ZENs on prepubertal gilts uterine endometrial glands. Explants were treated with 30 µM ZEN, ZEN-14-P, ZEN-16-P for 24 h. (A) Endometrial explants were stained with Hematoxylin phloxine saffron (HPS). and observed by bright-field microscope (B) Histogram shows the number of endometrial glands increased by each compound compared to the negative control. The reported values are the least square of means (LSM) and 95% confidence interval (CI) at least three independent replicates. Compared to the negative control, *p ≤ 0.05; **p ≤ 0.01. Compared with ZEN, ns, not significant.

### Biotransformation of ZEN and its phosphorylation products involves different metabolic paths

Considering the assessment of cell viability, oxidative stress, pro-inflammatory activity and estrogenic activity, C-14 or C-16 phosphorylation showed clearly to not reduce the toxicity of ZEN. Hence, the question arises whether the observed toxicities are intrinsic features of ZEN-14-P and ZEN-16-P, or they are due to conversion back to ZEN. Using LC-MS/MS, we traced metabolic pathways involved in the biotransformation of ZEN and its phosphorylation products in IPEC-J2 and Ishikawa cells. Figure 6 compares the phase I and phase II metabolite profiles reported following 24-hour incubation or IPEC-J2 and Ishikawa cells with ZEN and phosphorylated ZENs, while Figure 7 summarizes the putative metabolic pathways involved in the biotransformation each compound in the porcine intestinal cell line and the human endometrial cell line. The results indicate that under the applied conditions, phosphorylation products and ZEN converted into different metabolites through phase-I and phase-II biotransformation. For instance, in IPEC-J2 cells, although phosphorylated ZENs massively converted to ZEN, they did not produce the ZAN and α-ZAL reported when directly incubating ZEN. Likewise, ZAL-GLR produced from phase II biotransformation of ZEN in IPEC-J2 cells was not detected in the biotransformation of phosphorylated ZENs, while ZEN-14-GLR, ZOL-14-GLR and ZAN-S or ZOL-S were produced from phosphorylated ZENs, and not from ZEN.

**Figure 6.**
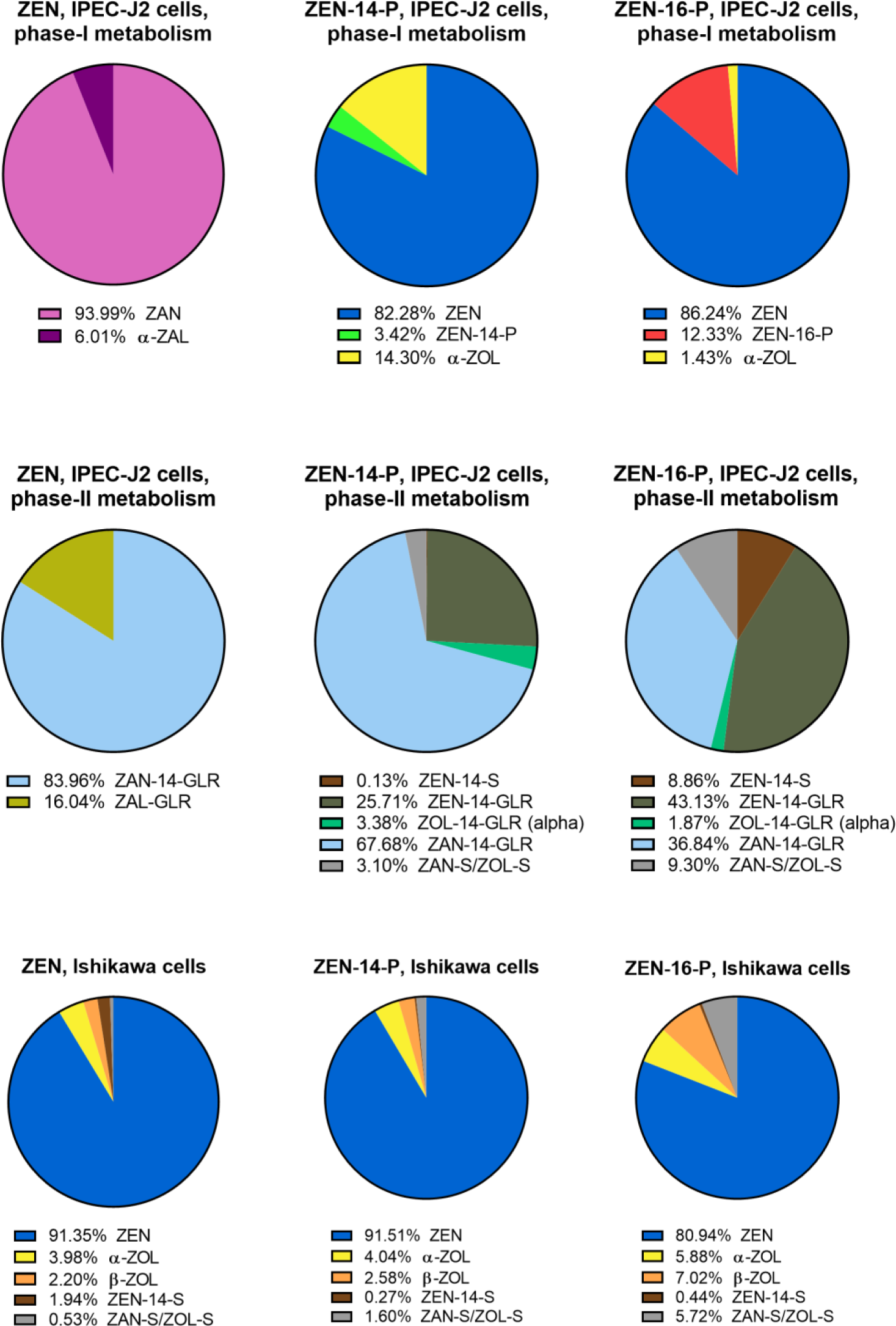
Phase-I and Phase-II metabolites of ZEN and C-14 and C-16 phosphorylation of ZENs after incubation with IPEC-J2 and Ishikawa cells.

**Figure 7.**
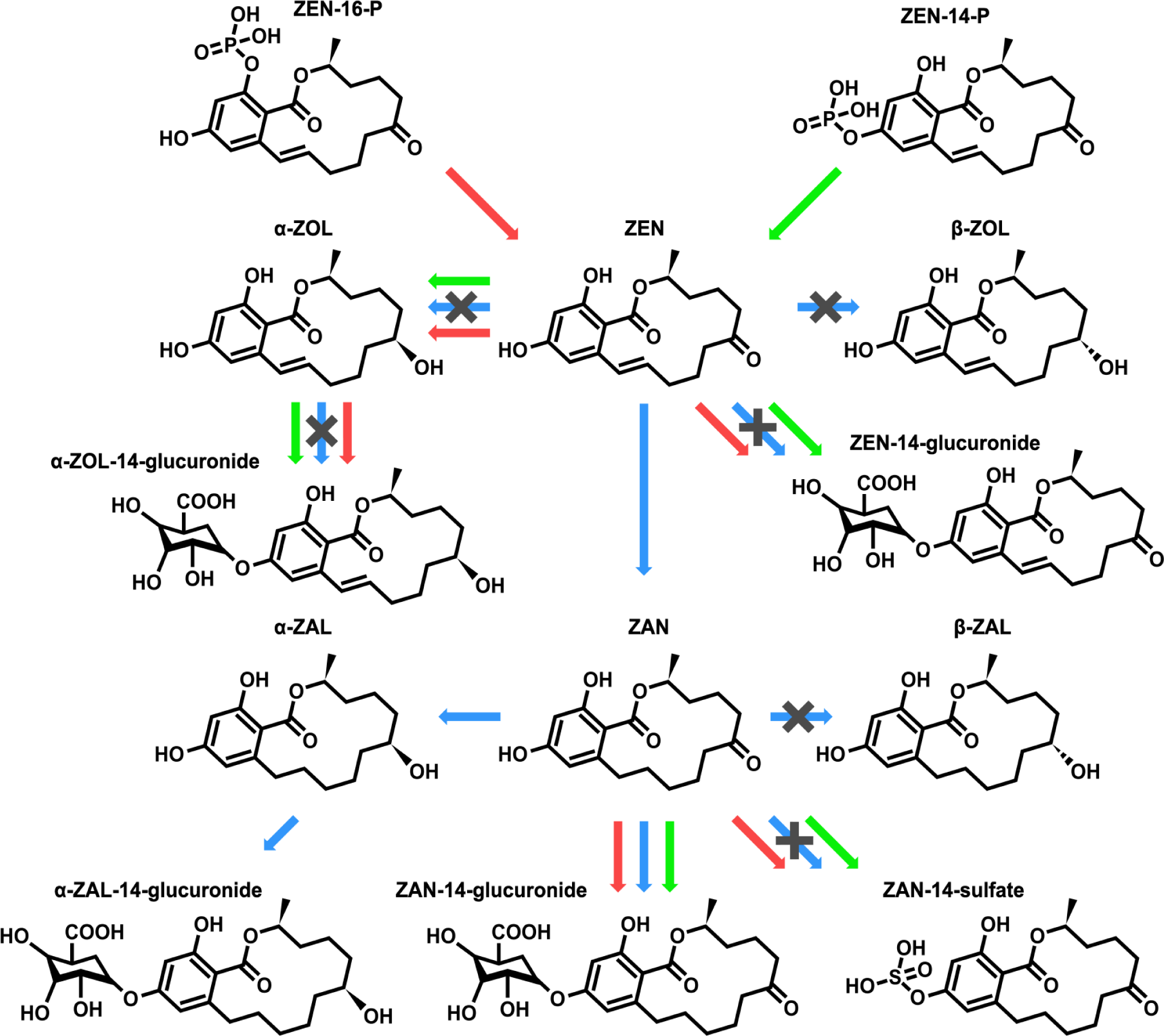
Biotransformation pathways of ZEN and C-14 and C-16 phosphorylation of ZENs

## DISCUSSION

ZEN causes significant human and animal health problems, resulting in substantial economic loss within the food industry and livestock farming.^2^ At the molecular level, ZEN binds to the estrogen receptors, generating an estrogen-like response, hence acting as an endocrine disruptor.^4^ At the cellular level, ZEN induces overproduction of reactive oxygen species, resulting ultimately in cell death.^20^ At the organ level, ZEN can alters the immune homeostasis, which in a bovine endometrium model, resulted in upregulation of the expression of proinflammatory cytokines, and hindering of the interaction between sperm and endometrial cells.^21^ In humans, exposure to ZEN has been associated with premature puberty.^22^ Therefore, effective detoxification methods should be developed to protect from the widespread presence and detrimental impact of ZEN and its metabolites in humans and animals. Two regioisomeric phosphorylated metabolites of ZEN, ZEN-14-phosphate and ZEN-16-phosphate, produced by *Bacillus* sp. S62-W, have been reported recently.^13^ So far, the main phase II metabolite that has been known from ZEN conjugation by detoxifying fungal and bacterial strains is ZEN-14-sulfate.^23–24^ Blockade of the phenolic hydroxyl group at C-14, in case of ZEN-14-sulfate, led to a substantial loss of estrogenicity, when compared to ZEN.^25^ Moreover, compare to ZEN, ZEN-14-sulfate demonstrated a milder effect on oxidative and thermal stress resistance of the model *Caenorhabditis elegans* organism.^26^ Since the toxicity of ZEN-14-P and ZEN-16-P compare to ZEN is still unknown, we used different biological models to assess the relevance of phosphorylation as a deactivation biotransformation path for ZEN.

Pigs exhibit significant sensitivity to ZEN and congeners.^9^ In our study, the porcine intestinal epithelial IPEC-J2 cell line that retains most of its original epithelial nature was selected as the intestine is the first barrier against food and feed contaminants and the first target for mycotoxins.^27^ Prepubertal gilt uterine explants were used to investigate the impact of phosphorylation on the estrogenic activity of ZEN. In a context of the 3Rs (replace–reduce–refine), the explant culture approach makes it possible to reduce the number of animals used while keeping enough strength for biological conclusion.^28^ Each of the six pigs used in this study allowed the investigation of all the five experimental conditions. The whole experiment (6 data-points × 5 experimental conditions) was achieved with only six animals, while the same statistical power would require more than 30 animals. Beside the porcine models, the human endometrial Ishikawa cells too were used to allow extrapolation of the findings in this study to other species. This cell line is sensitive to estrogenic stimulation.^29^

In this study, the cytotoxic effects of the two regioisomeric phosphorylation metabolites were first compared with ZEN using porcine IPEC-J2 cells and human endometrial Ishikawa cells. The results clearly indicate that phosphorylation does not decrease the cytotoxicity induced by ZEN, but rather increases it. Likewise, oxidative stress induction by ZEN in IPEC-J2 cells is not affected by C-14 or C-16 phosphorylation. These findings suggest that the phosphorylated metabolites, in line with ZEN, may activate the intrinsic and extrinsic apoptotic pathways.^30^ It is likely that these phosphorylated metabolites generate CYP-mediated quinones similar to the ones underlying the oxidative stress induced by ZEN.^31–32^

Next, the impact of phosphorylation on the pro-inflammatory activity of ZEN was analyzed in the porcine intestinal IPEC-J2 cells. Previous studies have reported pro-inflammatory cytokine release in porcine intestinal IPEC-J2 cells exposed to ZEN.^33–34^ In this study, the effects of two concentrations of ZEN, ZEN-14-P and ZEN-16-P on the expression of the pro-inflammatory cytokines *IL-1β*, *IL-6*, *IL-8* and *TNF-α* in IPEC-J2 cells were compared. At the lowest tested concentration, the phosphorylated derivatives displayed a pro-inflammatory activity similar level to ZEN, which suggests that the phosphorylated regioisomers retained the pro-inflammatory activity of ZEN. Interestingly at the highest concentration, the phosphorylated compounds induced lower upregulation of the cytokine expression, compare to ZEN. However, rather than an hypothetic quenching of the pro-inflammatory activity by the phosphorylation of ZEN, the global bell-shaped activation pattern of the cytokines observed for ZEN and phosphorylated derivatives as well, is consistent with the trigger of suppressors of cytokine signalling (SOCS). These proteins are induced upon stimulation by cytokines and inhibit the production of the same cytokine via a negative feedback mechanism.^35^ Mycotoxin-induced SOCS have been shown to regulate the signalling pathways mediated by the cytokine receptor superfamily members.^36^

The key biological feature of ZEN is its estrogenic effect on ^23, 37^ as it impairs estrogen signaling activity, disturbing reproductive hormones, and damaging organ histology.^38–39^ Therefore, this study investigated the estrogenic activity of ZEN-14-P and ZEN-16-P in comparison to ZEN, by observing alkaline phosphatase (ALP) activity and selective estrogen-responsive gene expression in Ishikawa cells, and the proliferation of endometrial glands using pre-pubertal gilt uterine explants. ALP activity is regulated by estrogens in an ER-dependent manner in Ishikawa cells and considered a sensitive means to assess ER agonists and antagonists.^29^ This was confirmed by our findings, in which ALP activity in Ishikawa cells was significantly stimulated by various concentrations of both ZEN and phosphorylated ZENs, just like the positive control E2. Moreover, the observed ALP inductions were suppressed in the presence of the ER high-affinity antagonist ICI 182 780, thus proving the ER-dependency of the estrogenic effects.

Estrogens have shown to mediate ALP gene expression in human endometrial cells by binding to specific estrogen receptors.^40^ Moreover, a distinct modulation of alkaline phosphatase isoenzymes has been reported when comparing the two estrogenic compounds 17β-estradiol and xanthohumol.^41^ Alkaline phosphatases exist in four distinct isoenzymes that play crucial roles in various significant functions, including hydrolysis of a large spectrum of phosphate-containing physiological compounds.^42^ ALPP and ALPG are the two isoenzymes that are expressed in endometrial tissue.^43^ To clarify the isoenzymes of alkaline phosphatase involved into the ALP activity detected for ZEN and for phosphorylation products, we investigated the expression of the placental alkaline phosphatase gene *ALPP*, and the germ cell alkaline phosphatase gene *ALPG* in Ishikawa cells exposed to either ZEN, or ZEN-14-P or ZEN-16-P. We also investigated upstream induction of alkaline phosphatase activation, the expression of potentially involved receptors, the progesterone receptor gene *PGR*, the estrogen receptor genes *ESR1* and *ESR2*, and the G protein-coupled estrogen receptor gene *GPER1.* Our results indicate that ZEN and phosphorylated compounds, activate the same ALP isoenzymes, and involve the same upstream receptors. The results also suggest that the presence of the phosphate moiety on C-14 or C-16 does not impede the flexible structure that makes ZEN to be able to bind mammalian ERs as strongly as E2.^44^ Interestingly the upregulation of ALP isoenzyme genes was also associated with increased expression of the membrane estrogen receptor *GPER1*. So far, the increase in ALP activity has only been associated with changes in the expression of nuclear estrogen receptors.^45^ The lack of difference in estrogenicity for ZEN and its phosphorylated conjugates observed *in vitro* in this study was paralleled by the results obtained on the *ex vivo* prepubertal gilt uterine explants model used as an alternative to the traditional rodent uterotrophic assay.^46^ Similar to E2, ZEN but also the phosphorylated ZENs induced the proliferation of endometrial glands after 24 h exposure. This rapid tissue proliferation is in line with the tumorigenesis associated with the activity of ALP isoenzymes.^47^ Alterations of the aromatic moiety of ZEN are believed to decrease its toxicity.^25^ However, the present study highlights that C-14 or C-16 phosphorylation does not decrease the toxicity of ZEN. Phosphorylation increased the cytotoxicity of ZEN. C-14 and C-16 phosphorylation products induced oxidative stress, and activated the expression of pro-inflammatory cytokines, as well as they triggered estrogenicity. Therefore, we were interested to know whether the observed toxicities are intrinsic features of ZEN-14-P and ZEN-16-P, or they are due to conversion back to ZEN. Through LC-MS/MS analysis, we investigated the biotransformation of ZEN and phosphorylated metabolites in IPEC J2 and Ishikawa cells. LC-MS/MS results revealed that though phosphorylated ZENs are partially converted into ZEN, they also produce different phase I and phase II metabolites. Especially, different glucuronide or sulfate metabolites were produced through phase-II metabolism from ZEN and phosphorylated compounds in both IPEC-J2 and Ishikawa cells. Beside the different relative potencies that have been established for ZEN and it congeners, their variable affinity for albumin may be pivotal contributors to their intrinsic toxicity.^48–49^

To conclude, this study investigated the residual toxicity of the two bacterial phosphorylation products of ZEN, ZEN-14-P and ZEN-16-P. The phosphorylated ZENs significantly decreased cell viability. Similar to ZEN, phosphorylation products induced significant oxidative stress, activated the expression of pro-inflammatory cytokines, and demonstrated estrogenic activity through upregulation of estrogen-responsive genes, activation of alkaline phosphatase and proliferation of endometrial glands. LC-MS/MS analysis pointed that although phosphorylated ZENs are partially hydrolyzed to ZEN, different metabolic pathways are involved in their biotransformation. Altogether, these results indicate that phosphorylation may not be a relevant biotransformation pathway for the detoxification of ZEN, and highlight the need to continue the search for suitable biological systems of ZEN neutralization in agricultural commodities.

## ABBREVIATIONS

ER: Estrogen receptor
GAPDH: Glyceraldehyde-3-phosphate dehydrogenase
RPL-32: Ribosomal Protein L32
IL-6: Interleukin-6
IL-8: Interleukin-8
IL-1β: Interleukin-1β
IPEC-J2: Intestinal porcine epithelial cells from jejunum
IPEC-1: Intestinal porcine epithelial cell line-1
LC-MS/MS: Liquid chromatography-tandem mass spectrometry
ROS: Reactive oxygen species
TNF-α: Tumor necrosis factor- α
α-ZAL: α-Zearalanol
α-ZOL: α-Zearalenol
β -ZAL: β -Zearalanone
Β-ZOL: β -Zearalenol
ZAN: Zearalanol
ZEN-14-GLR: Zearalenone-14-Glucuronide
ZEN-14-S: Zearalenne-14-Sulfate
ZOL-14-GLR: Zearalenol-14-Glucuronide
ZAN-14-GLR: Zearalanol-14-Glucuronide
ZAN-S: Zearalanol-Sulfate
ZOL-S: Zearalenol-Sulfate
ZAL-GLR: Zearalanone- Glucuronide

## ACKNOWLEDGMENTS

The author thanks Cai Goudong for his support and suggestions on this study

## DATA AVAILABILITY STATEMENT

All data pertinent to this study are contained in the article or available upon request. For all requests, please contact Dr. Imourana Alassane-Kpembi

## ETHICS STATEMENT

All animal experimentation procedures were reviewed and approved by the Animal Ethics Committee of the Faculty of Veterinary Medicine of the University of Montreal (Approval number #20-Rech-2071), and the pigs were cared for in accordance with the Canadian Council on Animal Care guidelines

